# Learning Continuous Potentials from smFRET

**DOI:** 10.1101/2022.09.12.507719

**Authors:** J. Shepard Bryan, Steve Pressé

## Abstract

Potential energy landscapes are useful models in describing events such as protein folding and binding. While single molecule fluorescence resonance energy transfer (smFRET) experiments encode information on continuous potentials for the system probed, including rarely visited barriers between putative potential minima, this information is rarely decoded from the data. This is because existing analysis methods often model smFRET output assuming, from the onset, that the system probed evolves in a discretized state-space to be analyzed within a Hidden Markov Model (HMM) paradigm. By contrast, here we infer continuous potentials from smFRET data without discretely approximating the state-space. We do so by operating within a Bayesian nonparametric paradigm by placing priors on the family of all possible potential curves. As our inference accounts for a number of required experimental features raising computational cost (such as incorporating discrete photon shot noise), the framework leverages a Structured-Kernel-Interpolation Gaussian Process prior to help curtail computational cost. We show that our Structured-Kernel-Interpolation Priors for Potential Energy Reconstruction from smFRET (SKIPPER-FRET) analysis accurately infers the potential energy landscape from a smFRET binding experiment. We then illustrate advantages of SKIPPER-FRET over standard HMM approaches by providing information, such as barrier heights and friction coefficients, otherwise inaccessible to HMMs.

**SIGNIFICANCE:** We introduce SKIPPER-FRET, a tool for inferring continuous potential energy landscapes, including barrier heights, from single molecule smFRET data. We benchmark on synthetic and experimental data.

## INTRODUCTION

Potential energy landscapes are useful continuous space model reductions employed across biophysics (1–6). For example, potentials can model dynamics along smooth reaction coordinates (3, 7–10) including the celebrated protein folding funnel (3, 8, 11). They also provide a natural language from which to calculate thermodynamic quantities (12–14). Furthermore, shapes of landscapes, including barrier heights, can provide insight into molecular function (15, 16) such as molecular motor dynamics (16). As such, inferring accurate potentials is a crucial step towards gaining insight into biophysical systems.

One way by which to decode potential energy landscapes from biological systems is through single molecule Fluorescence Resonance Energy Transfer (smFRET) experiments (17–19). Most commonly, smFRET works by tagging two locations of a biomolecule with pairs of fluorophores. When in proximity, the fluorophore excited by the laser (the donor) may transfer its excitation, via dipole-dipole coupling, over to the acceptor fluorophore (20). As the distance between the donor and acceptor fluorophores change, so too does the efficiency of dipole-dipole energy transfer resulting in higher donor emission rates when fluorophores are further apart. Conversely, more photons are emitted from the acceptor when fluorophores are in close proximity (20). As such, it is common to use the proportion of donor and acceptor photons counted in a given time window, the FRET efficiency, to estimate the pair fluorophore distance (17, 21).

To deduce energies from smFRET data it is common to immediately assume a discrete state-space and invoke Hidden Markov Models (HMMs) in the ensuing analysis (22–26). HMMs work by partitioning the observed smFRET efficiencies into discrete levels coinciding with distinct states. One can then use smFRET data to infer the number of states in addition to the associated transition rate parameters and pair distances (2, 25), which in turn can be used to infer the potential energy of the states using the Boltzmann distribution (2, 27).

The above approach is useful in gaining quantitative insight into systems well approximated by discrete states (10, 23, 24, 26). However the above formulation is not appropriate when the dynamics occur along a continuous reaction coordinate poorly approximated by well separated discrete-states (5, 11).

Furthermore, while HMMs can be used to infer each state’s relative energies, they cannot reveal energy barriers between states without preexisting knowledge of internal system parameters, such as the landscape curvature and internal friction, due to loss of information inherent to the discretization process (2, 28). The inability to infer accurate potential energy barriers from a single data set without the knowledge of hidden internal parameters is a major limitation of HMMs applied to smFRET data.

As such, a method capable of inferring potential energy landscapes, including barrier heights, along a continuous coordinate would greatly enhance the resolution with which we can probe biophysical systems and lend deeper insight into protein folding (11), protein binding (3), and the physics of molecular motors (16).

Here, we develop a method to decode a continuous potential from smFRET data without resorting to discrete state-space assumptions inherent to HMM modeling. We do so by incorporating a detailed, physics-informed likelihood distribution describing the relationship between measurements and a potential energy landscape. We then infer the most probable potential energy landscape within the Bayesian nonparametric paradigm by placing a prior on the potential energy landscape with support over the family of all putative continuous curves. Our prior distribution is built upon the Structured-Kernel-Interpolation Gaussian-Process (29), which allows for inference of continuous potentials while simultaneously avoiding the costly cubic scaling of conventional Gaussian-Process regression. Cubic scaling becomes especially problematic as we insist on incorporating realistic measurement features into our likelihood.

We show that our Structure-Kernel-Interpolation Priors for Potential Energy Reconstruction from smFRET (SKIPPER-FRET) analysis unveils the full potential energy landscape, including barrier heights and friction coefficients within reasonable computational time. The essence of SKIPPER-FRET is described in cartoon form in figure 1. We benchmark SKIPPER-FRET on synthetic/simulated data as well as experimental data.

**Figure 1:**
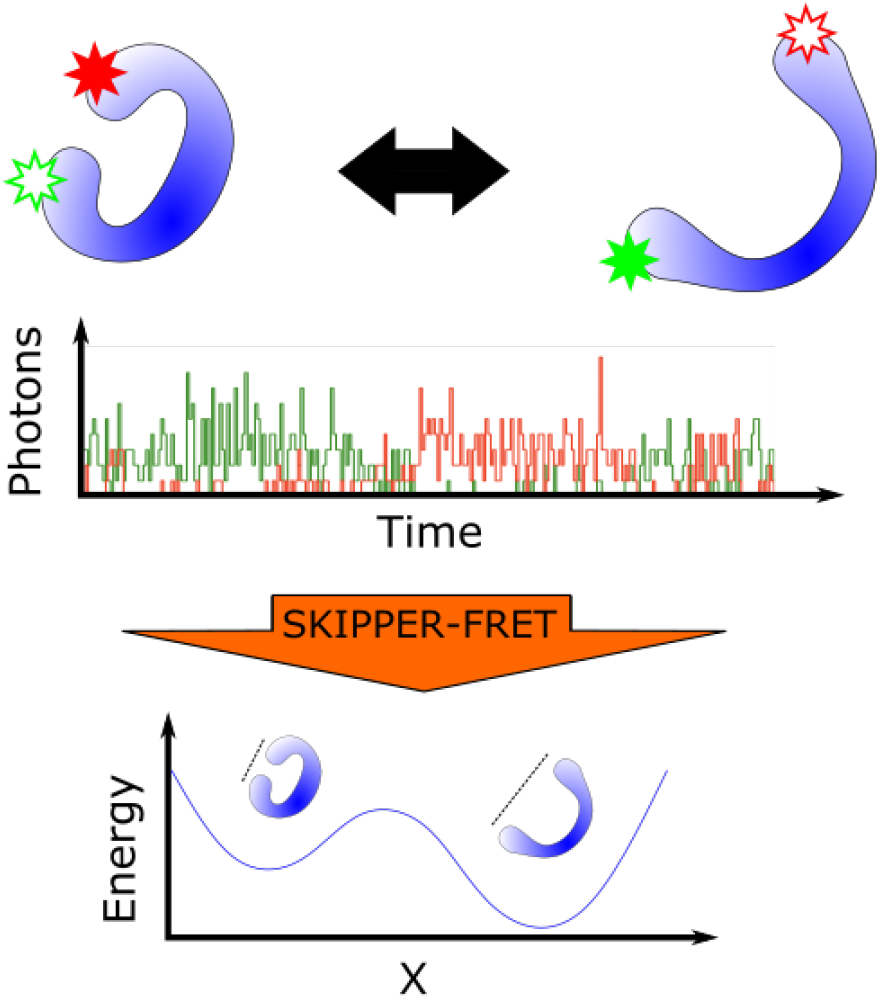
Cartoon description of SKIPPER-FRET. At the top we see a protein switching between two conformations over time. The protein is labeled with a donor and acceptor fluorophore. As the protein changes configuration the FRET efficiency between the fluorophores also changes. In the middle panel we illustrate a typical trace containing the number of red and green photons over time. On the bottom panel we show the outcome of SKIPPER-FRET analysis used to infer the potential energy landscape along the reaction coordinate probed.

## METHODS

Our goal is to learn potentials given photon arrival data from two channels assuming continuous illumination smFRET data. Generalization to pulsed data is possible and addressed in the Discussion 0.4. In this section, we present a forward model describing how a potential gives rise to the data we collect. Next, we describe an inverse model allowing us to infer the potential directly from the data along with a numerical algorithm developed to sample from our high dimensional posterior. We conclude by summarizing the experiment we use to validate our method.

### 0.1 Forward model

In our framework, we imagine placing donor and acceptor fluorophores at two points whose relative distance varies with time. That is, we envision either monitoring a molecule undergoing configurational changes along a reaction coordinate or a pair of molecules binding and unbinding; in either case, our formulation is identical. The dynamics of the probes with respect to each other are dictated by a potential we wish to deduce. The labeled system is exposed to continuous illumination in which both fluorophores will be excited. Donor excitations have a position dependent probability of FRET transfer, whereas acceptor excitations are treated as a source of background. We describe this process in detail in this section.

#### 0.1.1 Pair distance

We begin by assuming that the distance of interest evolves according to Langevin dynamics (5, 27)

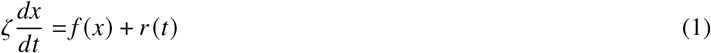

with unknown *ζ* (the friction coefficient) and unknown spatially dependent force (*f* = −∇*U*). In the above, *r* (*t*) is the thermal noise whose moments are

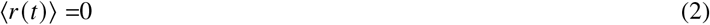

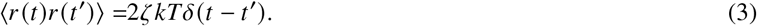

Here *kT* is the usual thermal energy and ⟨… ⟩ denotes an average over thermal noise realizations.

Under the Ito approximation (30), we can evaluate equation 1 on a fine grid of time levels,

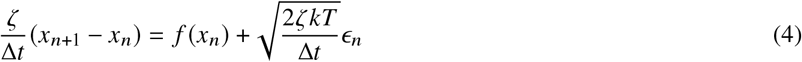

where *x*_*n*_ is the distance at time level *n*, Δ*t* is the time step size, and *ϵ*_*n*_ is a normally distributed random variable with mean 0 and variance 1. We can rewrite the probability of *x*_*n*+1_ as follows

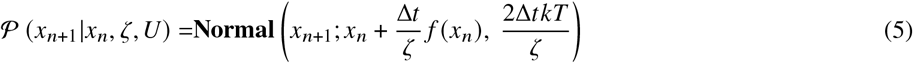

which reads “the probability of *x*_*n*+1_ given *ζ, U* and the previous position (*x*_*n*_) is a Normal distribution with mean 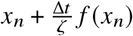 and variance 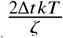”. Here we let *N* be the number of time levels and let ***x***_1:*N*_ represent the set of all positions at those time levels. Note that the time step, Δ*t*, must be chosen to be small enough that the Ito approximation be valid but, in principle, need not coincide with the measurement time scale.

An important note is in order. When analyzing data from binding experiments, we envision a donor tagged immobilized biomolecule interacting with an acceptor tagged binding agent. In this setup we interpret the pair distance, *x*, as the distance between the donor fluorophore and the nearest acceptor fluorophore with the understanding that the identity of the acceptor fluorophore may change over time.

#### 0.1.2 Photon measurements

To model photon counts, we make a number of physically reasonable assumptions. First, we assume that time scales over which pair distances vary are much slower than fluorophore excited state relaxation times (21) (microseconds or slower versus nanoseconds (2, 10)). Secondly, we assume that the small absorption cross section of the fluorophores results in a low excitation rate compared with the relaxation rate. Thus, the interphoton arrival time is dominated by the excitation rate, *λ*_*X*_ (21).

As the pair distance is assumed to remain constant over the whole time step (see equation 5), the FRET rate will also be assumed constant (with changes approximated as occurring when time levels change). Thus photon arrival times and the order of photon colors within a time step provide no additional information. In this regime, the probability of the number of measured green, *g*_*n*_, and red, *r*_*n*_, photons are drawn from a Poisson distribution (see SI section S0.5)

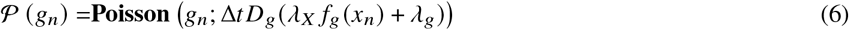

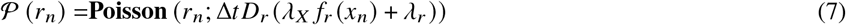

where *λ*_*X*_ is the donor excitation rate, *λ*_*g*_ is the green photon background rate, *λ*_*r*_ is the red photon background rate (which includes the direct acceptor excitation rate), *D*_*g*_ and *D*_*r*_ are detector efficiencies, and *f*_*g*_ (*x*_*n*_) and *f*_*r*_ (*x*_*n*_) are the fraction of photons emitted by the FRET pair detected in the green and red channel, respectively, calculated from the FRET rate as a function of position, FRET (*x*). The crosstalk matrix, which encodes the rate at which a red photon is measured to be green and vice versa, reads as follows

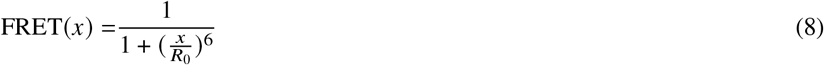

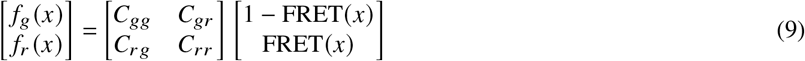

where *R*_0_ is the characteristic distance for the acceptor donor pair at which the FRET efficiency is 0.5 and *C*_*i j*_ is the probability that a photon with color *i* is detected by detector *j*. For example, *C*_*rg*_ is the probability that a red photon is detected by the green photon detector.

### 0.2 Inverse model

Our goal is the create a probability distribution for the potential energy landscape, *U*(*x*), the pair distance trajectory, ***x***_1:*N*_, the excitation rate, *λ*_*X*_, the background photon rates, *ζ*_*r*_ and *ζ*_*g*_, and the friction coefficient, *ζ*, given a series of photon measurements, ***g***_1:*N*_ and ***r***_1:*N*_. Note that detector efficiencies, *D*_*g*_ and *D*_*r*_, and the crosstalk matrix can be calibrated separately and therefore do not need to be inferred. Using Bayes’ theorem we can express

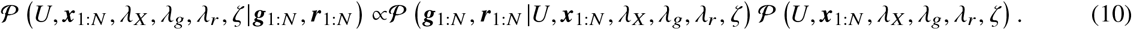

The first term on the right side of equation 10 is called the likelihood and is equal to the product of equations 6 and 7 for each time level. The second term is called the prior and can further be decomposed as follows

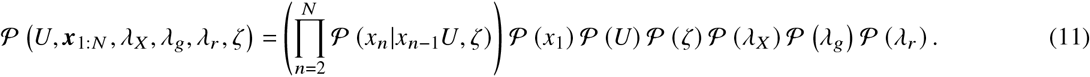

The first term on the right hand side, *𝒫* (*x*_*n*_ |*x*_*n*−1_*U, ζ*), is informed by dynamics model (equation 5). We are free to choose the remaining priors over *𝒫* (*x*_1_), *𝒫* (*U*), *𝒫* (*ζ*), *𝒫* (*λ*_*X*_), *𝒫*, (*λ*_*g*_)and *𝒫* (*λ*_*r*_).

We start by placing priors on our photon rates and friction coefficient. We know that our excitation rate, *λ*_*X*_, is strictly positive and, as such, an acceptable choice of prior is the Gamma distribution which has nonzero probability density along the positive real line

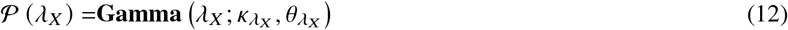

where 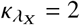 is chosen to make the mode of the distribution diffuse (i.e., create an uninformative prior), and 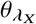 is chosen so as to give a mean expected value close to the average number of observed photons per frame. Similarly we set a Gamma prior on our background photon rates

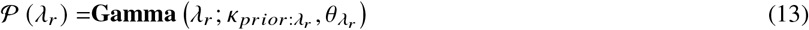

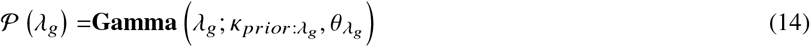

where we again choose 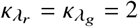 and choose values for 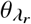 and 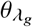 that give mean values close to the measured background rates. Similarly, because *ζ* is strictly positive, the Gamma distribution is also a good choice. As a prior over the friction coefficient, we choose

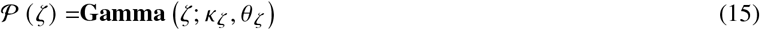

where *κ*_*ζ*_ = 2 and *θ*_*ζ*_ = 5000ag/ns are chosen to be minimally informative and allow some density over several orders of magnitude over the real positive line. Note that 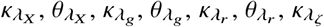, and 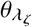 are hyperparameters whose exact values bear little weight on the final form of the posterior as more data are acquired (22)

Next we place a prior on our initial position. That is, under our dynamics model, equation (5), all positions, ***x***_2:*N*_, are directly conditioned on the previous position, i.e., the dynamics follow a Markov chain. As such, we must only place a prior on the position at the first time level, *x*_1_. For computational reasons alone, we choose a Normal distribution as the prior over *x*_1_ as it matches the form of the transition probability of equation 5,

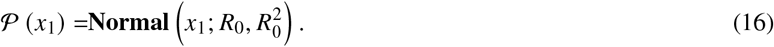

As the initial position is known to be around the characteristic FRET distance up to some uncertainty, we conveniently choose to center our distribution at *R*_0_ with standard deviation *R*_0_. The latter choices are immaterial in the presence of sufficient data.

Of greatest importance is our choice of prior on potential energy landscape, *U* (*x*). One natural prior choice is the Gaussian process (29, 31, 32) allowing us to sample from all putative curves without pre-specifying any functional form. However, a naive implementation of the Gaussian process is computationally intractable for large data sets as computational complexity scales cubically with the size of the data (29, 33). This is especially challenging given the lack of conjugacy between the likelihood and prior rendering direct sampling of the posterior infeasible.

Instead, we develop a computationally efficient adaptation of the Gaussian process leveraging recent advances in structured-kernel-interpolation Gaussian processes (SKI-GP) (29, 31). Briefly, SKI-GPs work by selecting a set of *M* nodes, 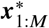, termed inducing points where we wish to exactly evaluate the potential. The value of the potential at the inducing points is itself drawn from a zero mean multivariate Normal distribution with some pre-specified covariance matrix

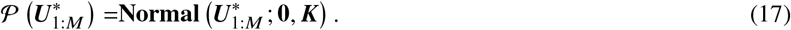

We then interpolate the value of the potential elsewhere (31). For example, collecting the force evaluated along the trajectory into a vector, *f*_1:*N*_, and the potential evaluated at the inducing points into a vector, 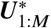, we can interpolate

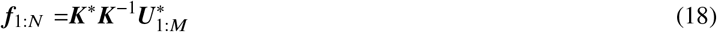

where ***K***^∗^ is the kernel matrix between the force at the trajectory and the potential at the inducing points inducing points with elements 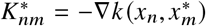 and ***K*** is the kernel matrix between the potential at the inducing points with elements 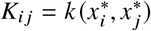.

We may then obtain the force at each position in the trajectory by taking discrete analog of the derivative of equation 18, *f* (*x*_*n*_) = −**∇***U*_*n*_.

Putting together all distributions and priors of our model, we attain a posterior for SKIPPER-FRET given by

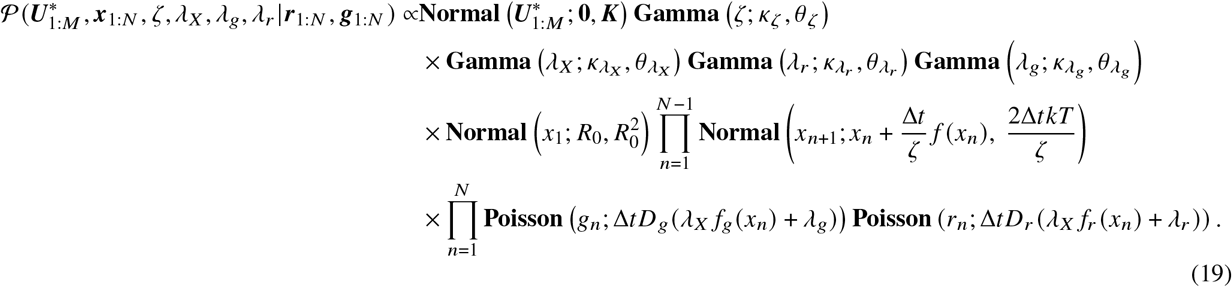

A graphical model of our full posterior, illustrating the conditional dependence of all variables, is shown in figure 2.

**Figure 2:**
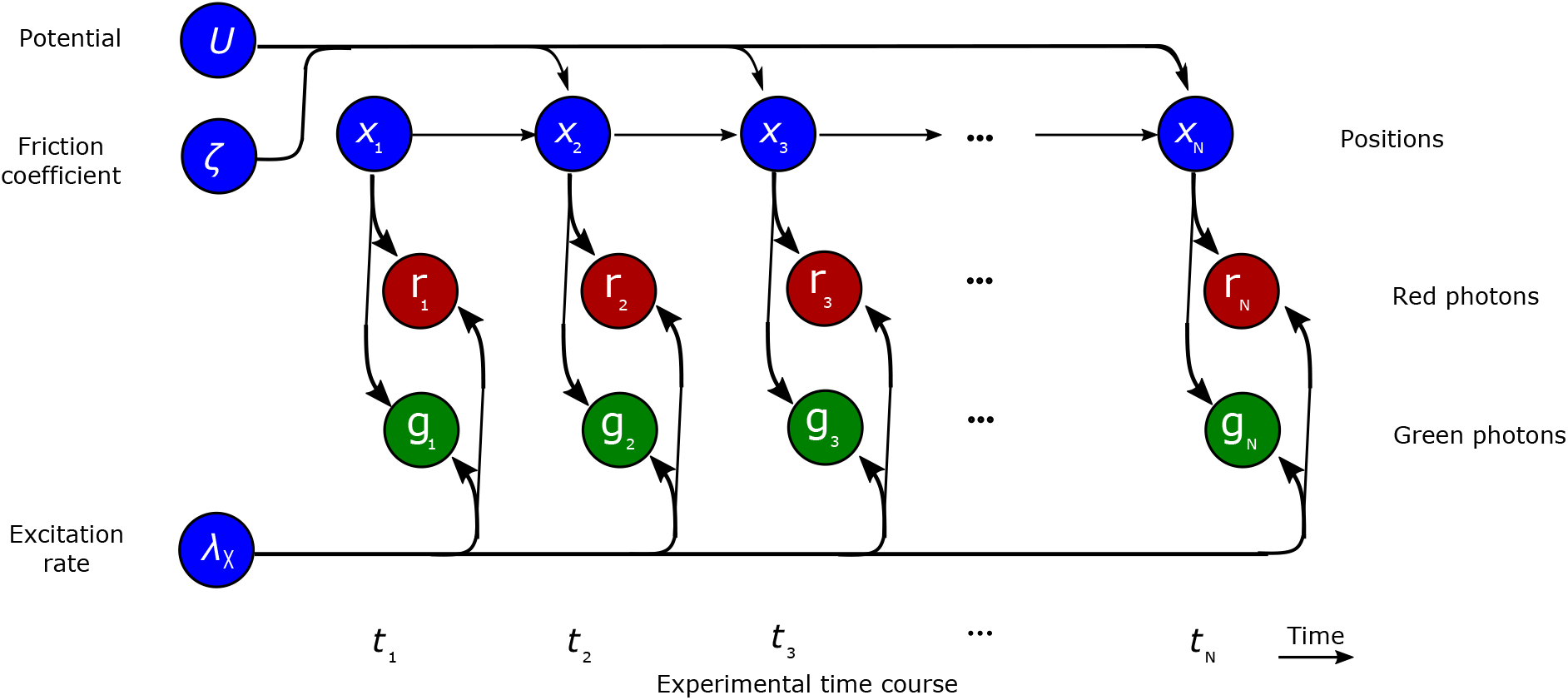
Graphical description of the model. Nodes (circles) represent random variables of our model and arrows connecting the nodes highlight conditional dependency. Blue nodes represent variables we wish to learn in our inference scheme, the red and green nodes represent the measured photon counts for each bin.

### 0.3 Algorithm

Our inverse model leaves us with a high dimensional posterior, equation (19), which does not attain an analytical form and cannot be directly sampled. Thus we propose to use a Gibbs sampling (22) scheme to draw samples from our posterior.

Briefly, Gibbs sampling works by starting from an initial guess for the parameters, then iteratively sampling each variable while holding other variables fixed. This scheme, where superscripts indicate the iteration index, is outlined below:

- **Step 1:** Start with an initial guess for each variable: 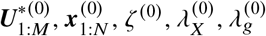, and 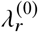.
- **Step 2:** For many iterations *i*,
  – Sample 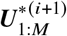 from 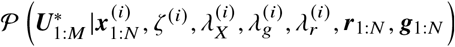.
  – Sample 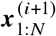 from 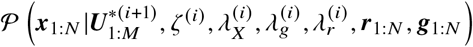.
  – Sample *ζ*^(*i*+1)^ from 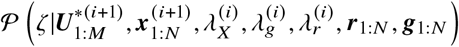.
  – Sample 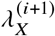 from 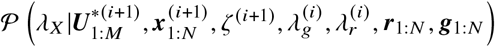.
  – Sample 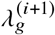 from 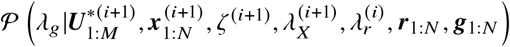.
  – Sample 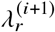 from 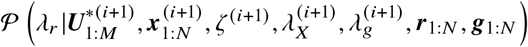.

The conditional probabilities appearing in Step 2 above are derived in SI section S0.6. Once sufficient samples have been generated (after burn-in is discarded (22)), we can use the sample average to provide point estimates for each variable or plot the distribution of all samples drawn.

### 0.4 Data acquisition

We analyze single photon smFRET data taken from an experiment probing the binding between the nuclear-coactivator binding domain (NCBD) of the CBP/p300 transcription factor and the activation domain of SRC-3 (ACTR) (34). ACTR and NCBD are both intrinsically disordered proteins (34–36). In the experiment, ACTR is surface immobilized and labeled with a donor dye (Cy3B). A solution including acceptor (CF660R) labeled NCBD is added. To probe the binding coordinate, we collect donor and acceptor photons as the NCBD binds and unbinds to ACTR. Further details on the data acquisition can be found in Zosel *et al* (34). Our analysis reveals the binding energy landscape of the ACTR-NCBD complex.

## RESULTS

In this section, we demonstrate our method on simulated and experimental data. We first show that our method can accurately infer the potential energy landscape from simulated smFRET data. We then demonstrate our method on real data from an experiment probing the binding energy landscape between NCBD and ACTR. We compare SKIPPER-FRET results to results obtained using a two state HMM that uses the same likelihood model as SKIPPER-FRET (see SI S0.10). We test the robustness of our method with respect to the number of data points in the SI.

We first analyzed simulated data using a simple double-well potential energy landscape. Figure 3 shows the data and the trajectory we infer. Figure 4 shows the SKIPPER-FRET potential energy landscape, the ground truth potential energy landscape, and the state energies inferred using a Bayesian HMM. The HMM does not infer full potential energy landscapes but rather just the energy and the pair distance of each state. As such, we cannot plot a full potential landscape for the HMM results and instead plot point estimates, with uncertainties, indicating the pair distance and energy levels of each state. We note that in order to compare our method against the ground truth in figure 4, we must define a common zero-point energy. Since only potential energy differences (not absolute values) are physical, the reference can be chosen arbitrarily. For our first data set, we chose the zero point energy to be the top of the barrier between the wells.

**Figure 3:**
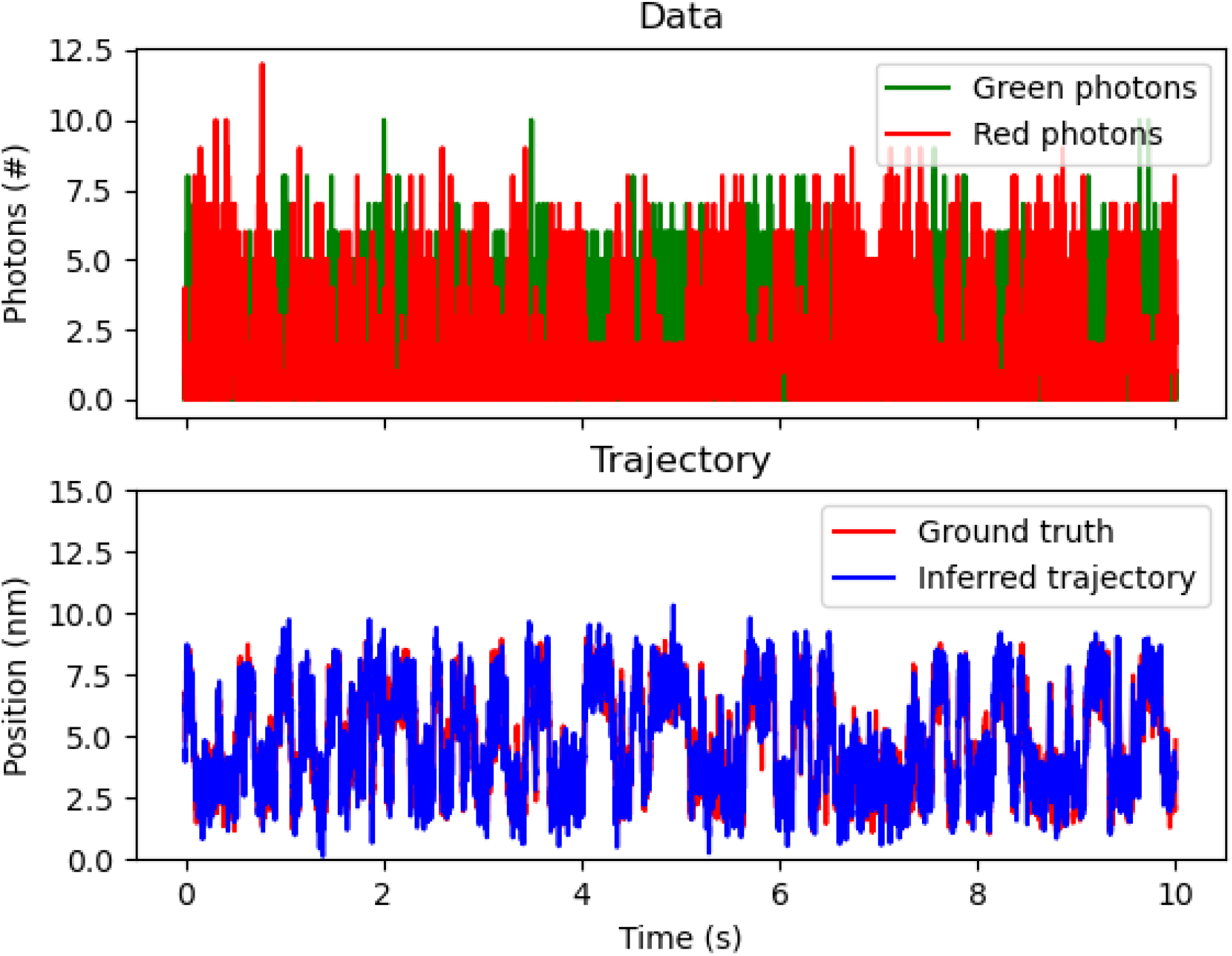
Demonstration on simulated data. Here we demonstrate our method on simulated data. (A) shows the raw data from the experiment including red and green photon counts binned every millisecond. (B) shows the inferred pair distance trajectory (blue) with the ground truth pair distance trajectory (red).

**Figure 4:**
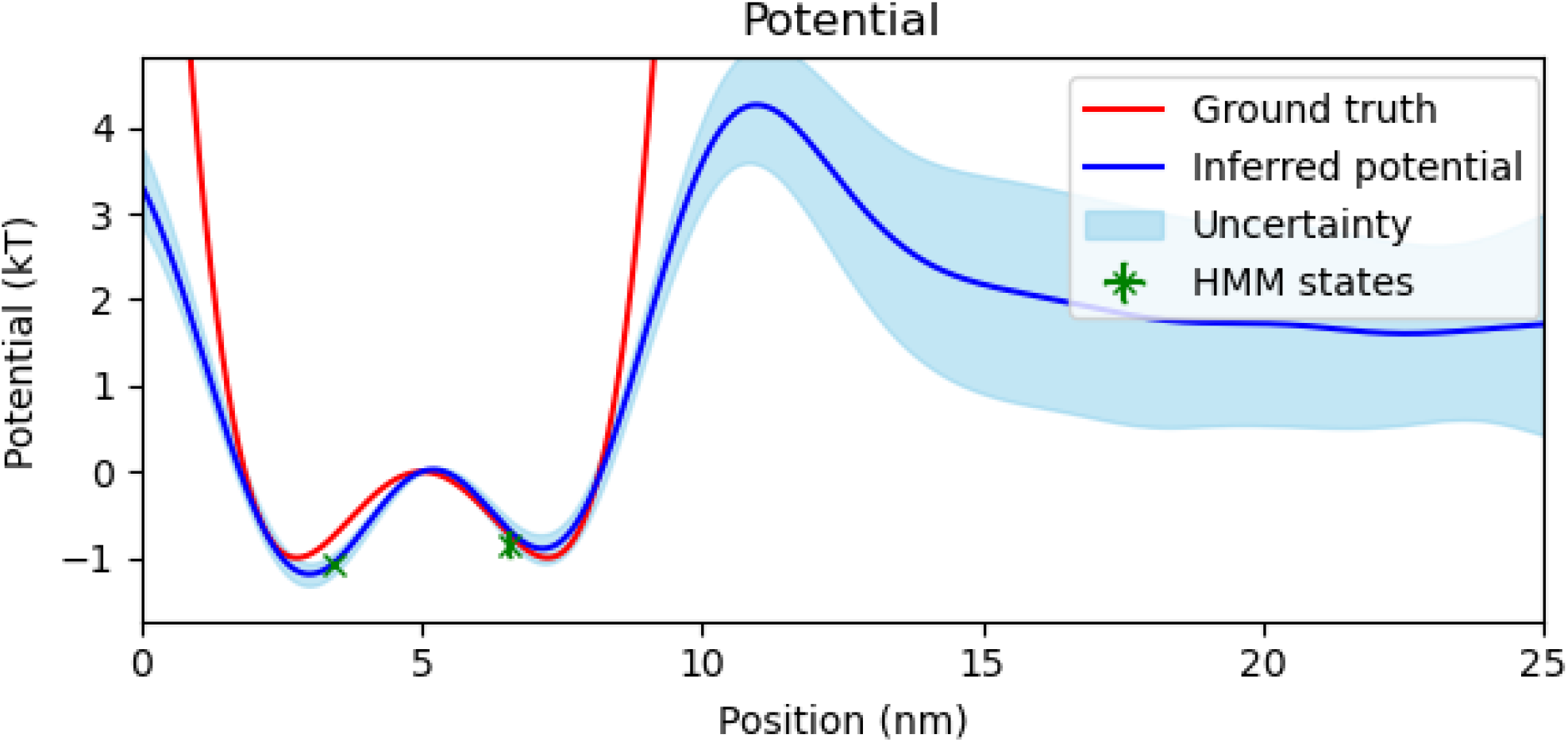
Simulated potential energy landscape. We show our inferred potential energy landscape (blue) with uncertainty (light blue) against the ground truth potential energy landscape used in the simulation (red). We additionally plot markers, with uncertainty, indicating the inferred state energy and pair distance using the HMM method (green). The common zero point energy was set at the top of the barrier at 5nm.

As seen in figure 3 the SKIPPER-FRET inferred pair distance trajectories are consistent with the ground truth trajectory. To be more quantitative, we note in figure 4 that our inferred potential energy landscape well minima and barrier height locations are all within 0.2nm of the ground truth. The inferred well energies were accurate within 0.1*kT* (−1.2 ± 0.1*kT* and −0.9 ± 0.1*kT* vs −1*kT* and −1*kT*). We note that as we move to the right of the rightmost well, the data becomes increasingly sparser and, consequently, our method returns the flat prior as the potential with high uncertainty.

We further note from figure 4 that while the energies inferred using a Bayesian HMM match the energies learned using SKIPPER-FRET, the pair distances inferred using the HMM deviate from both the ground truth and SKIPPER-FRET well minima. This is on account of the fact that the HMM ascribes a single specified pair distance to what is, in reality, a continuous range of pair distances near potential well minima. To estimate a single specified pair distance, the HMM finds itself effectively averaging the FRET efficiencies over those portions of the trajectory it deems as belonging to one state. This effective pair distance averaging is further complicated when the pair distance trajectory crosses a barrier in which case the HMM must somehow ascribe the dynamics when surmounting the barrier, which it cannot model, to one of the states.

Next we analyzed simulated data from a double-well potential where the far rightmost well is centered beyond the range of traditional smFRET measurements (at distance *>* 2*R*_0_ where less than 2% of absorbed photons are transferred to the acceptor). Such a potential mimics the data that we expect to see from the binding experiments we later analyze. Figure 5 shows the results where the zero point energy is set at the bottom of the leftmost well. As seen in figure 5, our method is able to infer the shape of the left well (where most photons are collected) and still manages to deduce, albeit with reduced accuracy, the shape of the barrier and the far well. The ground truth potential is enclosed within the uncertainty regions (one standard deviation) of our estimates at almost every point along the left well. Our method further infers a barrier height of about 2.5kT which is within 0.5 kT of the ground truth barrier height (2.9kT). On the right side of the barrier where the FRET efficiency drops dramatically, and we therefore have less information to inform the shape of the potential inferred, our estimate deviates from the ground truth with a correspondingly growing uncertainty. When comparing to the HMM method, we again see that the HMM method and SKIPPER-FRET estimate similar energies, but different well locations.

**Figure 5:**
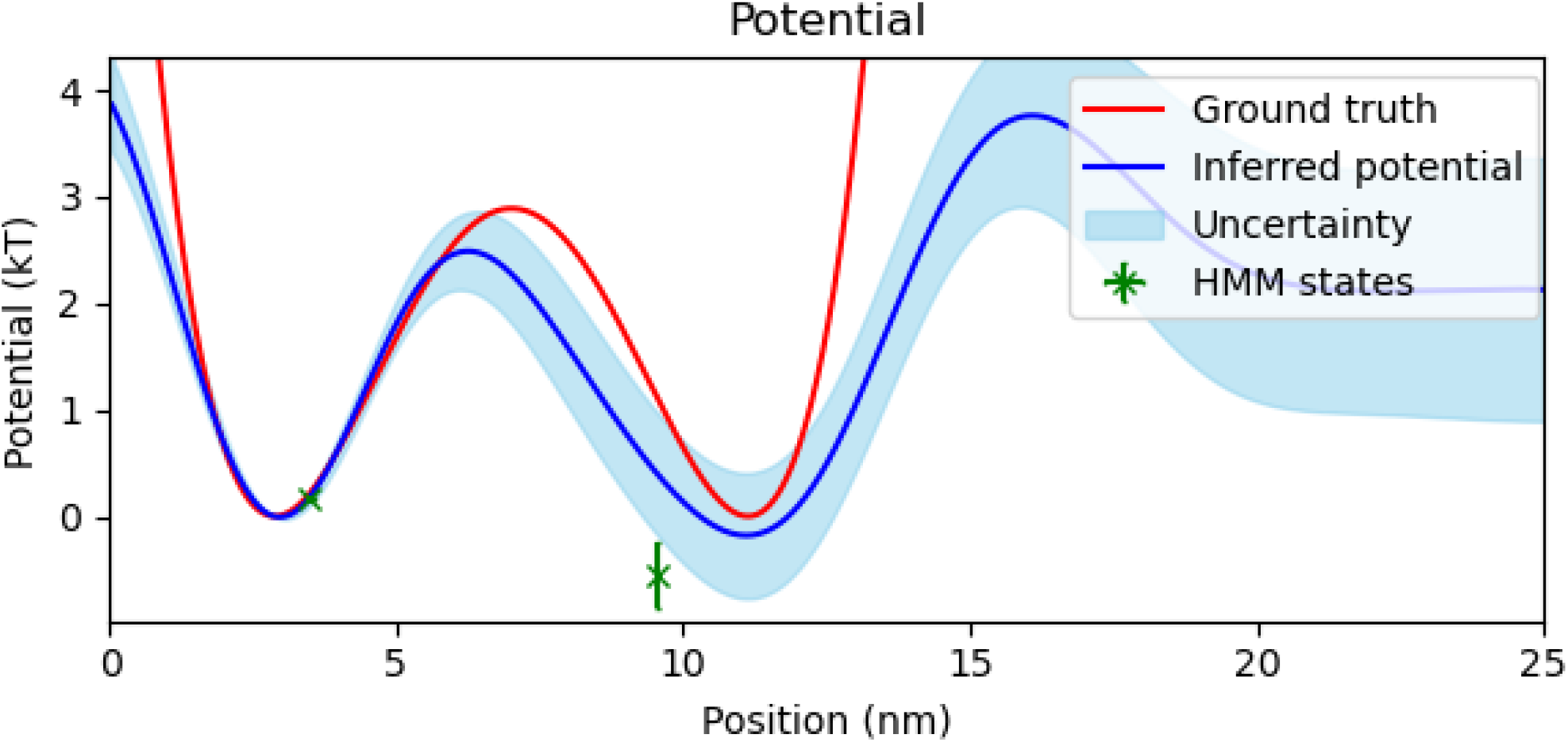
Simulated potential energy landscape when one barrier is far from the characteristic FRET distance. Here we analyze data simulated using an energy landscape in which one of the wells is outside of the characteristic FRET range. We compare our inferred potential energy landscape to the potential energy landscape inferred using the Bayesian HMM as well as the ground truth. We show our inferred potential energy landscape (blue) with uncertainty (light blue) against the ground truth potential energy landscape used in the simulation (red). We additionally plot markers, with uncertainty, indicating the inferred state energy and pair distance using the HMM method (green). The common zero point energy was set at the bottom of the leftmost well at 2.87nm

After successfully testing SKIPPER-FRET on simulated data, we now move on to the analysis of experimental data. In Figure 6, we show the inferred trajectory by applying our method to data from ACTR-NCBD binding-unbinding experiments (34). Based on independent analysis (35, 37, 38), we expect to find two states corresponding to a bound and an unbound state. Furthermore, looking at the raw data in figure 6 A, we immediately notice that there are alternating sections of high and low FRET efficiency in what appears to be two states. The corresponding inferred pair distance trajectory, as seen in figure 6 B, also alternates between two levels as expected.

**Figure 6:**
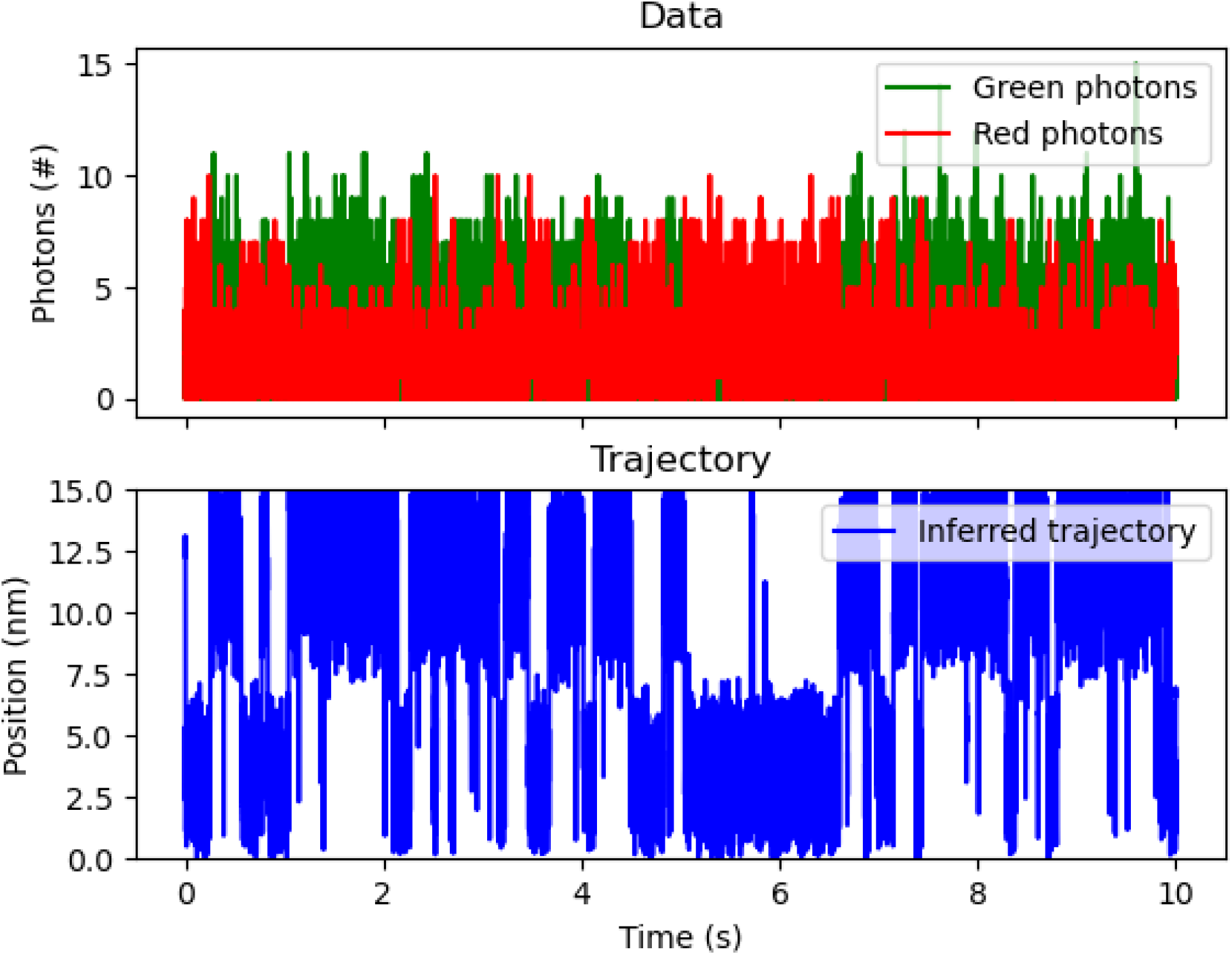
Demonstration on NCBD-ACTR. Here we demonstrate our method on data probing the energy landscape of NCBD-ACTR binding. (A) shows the raw data from the experiment including red and green photon counts. (B) shows the inferred pair distance trajectory (blue).

We also show the inferred potential energy landscape in Figure 7. Indeed, as expected, we see a double well shape. The left well in figure 7 can be interpreted as the binding energy between ACTR and NCBD, while the right well can be interpreted as the chemical potential energy required to remove NCBD from a volume surrounding the ACTR.

**Figure 7:**
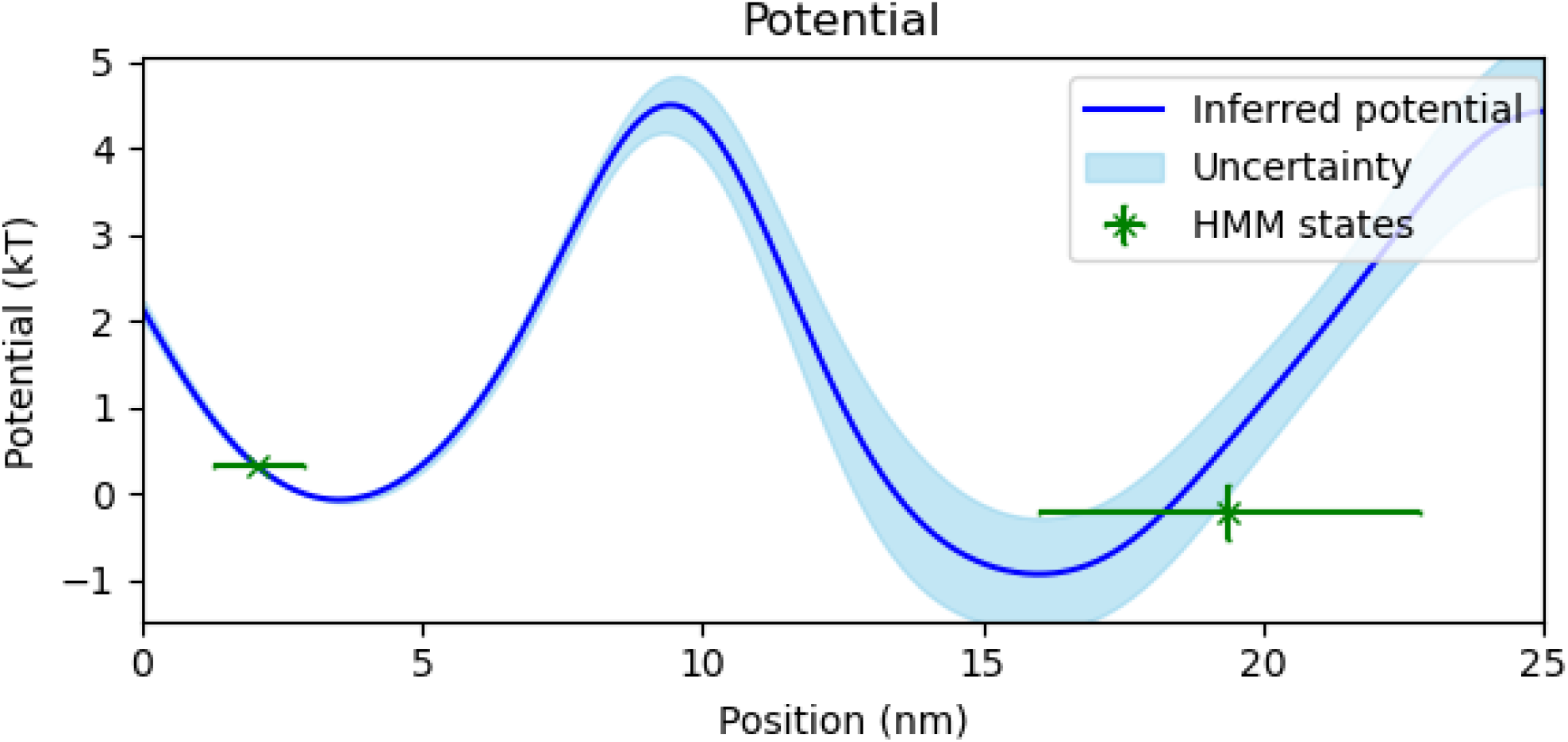
NCBD-ACTR potential energy landscape. Here we compare our inferred potential energy landscape to the relative potential energy landscape inferred using standard HMM methods. We show our inferred potential energy landscape (blue) with uncertainty (light blue). We additionally plot markers, with uncertainty, indicating the inferred state energy and pair distance using the HMM method (green).

As the true energy landscape for ACTR-NCBD binding is unknown, we compare our results to the energy landscape inferred using a two state Bayesian HMM model with the same likelihood model as SKIPPER-FRET (see SI S0.10). As seen in figure 7, the energies inferred using the HMM method fall within our uncertainty regions, but position of the wells inferred using SKIPPER-FRET differ from those inferred using the HMM method. As explained earlier, this arises because, fundamentally, the HMM attempts to reconcile its discrete state picture with the Langevin model’s continuous formulation. As the HMM method does not provide barrier height, we cannot naturally accommodate the barrier inferred using SKIPPER-FRET without additional information (see SI 40).

## DISCUSSION

Inferring accurate potential energy landscapes is a critical step toward unraveling key biophysical phenomena including protein folding (11), binding (3, 8), and the dynamics of molecular motors (16). Here, we have developed a method orthogonal to the HMM paradigm to include continuous states which also yields barrier heights. We benchmarked our method on simulated and experimental data.

We showed that, if warranted, we can avoid making the discrete state assumption inherent to HMMs while the HMM only has access to energy barriers between states if we supply it with preexisting knowledge of the internal parameters of the reaction coordinate or if there are at least two data sets taken at different temperatures (see SI section 40). This is despite any single data set already encoding this information.

Key to our inference algorithm is the structured kernel interpolation Gaussian process (SKI-GP), which allows us to sample the potential energy landscape from a prior over all continuous curves while avoiding the costly cubic scaling requirements of a standard Gaussian process. Specifically, with the SKI-GP prior, we are able to distinguish the inducing point locations, 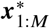, from the trajectory, ***x***_1:*N*_, so we do not have to calculate a new covariance matrix, ***K***, and its inverse, ***K***^−1^ at each iteration of our Gibbs sampler, thus saving us considerable computation time. This would not be possible using standard Gaussian process techniques.

Moving forward, there are ways in which we may improve SKIPPER-FRET. Firstly, our method, as it stands, deals with smFRET data from continuously illuminated sources. However, many smFRET experiments work using pulsed excitation (21, 39). We could modify our measurement model (equation (6)-equation (7)) to accommodate pulsed illumination by swapping the Poisson distribution, which assumes exponential waiting times between excitation, for a Binomial distribution, compatible with fixed window excitations.

Also, our method deals with dynamics along a single reaction coordinate assumed to be equivalent to the FRET pair distance. However, one can imagine situations in which the system’s dynamics are probed along an axis partly orthogonal to the FRET pair distance (40, 41) in a multi-dimensional incarnation of FRET with, say, one donor and multiple acceptor labels. For example, even in the case of ACTR binding to NCBD, analyzed in this manuscript, the ACTR may rotate with respect to the NCBD during binding. Cases with multiple degrees of freedom are traditionally studied using multicolor smFRET (41–44) or by pairing data analysis with molecular dynamics simulations (41, 45). In principle, one could use SKIPPER-FRET to infer potentials along degrees of freedom orthogonal to the FRET distance by including some mapping from the desired degree of freedom to the FRET pair distance in equation (8). As the FRET pair distance is often not directly tied to the reaction coordinate (39, 46) this may be a promising direction for future work.

Along these same lines, while our focus has, so far, been on learning one-dimensional potentials and demonstrating that we can learn barriers and potential shapes, avoiding the costly cubic scaling of standard GPs is also critical in deducing higher dimensional potentials. For instance, a HMM may, for example, distinguish between a fully connected and linear three-state model. Here, our one-dimensional reduction would need to be augmented to two dimensions in order for us to deduce these types of higher-dimensional features. Deducing features, such as potential ridges and valleys, in higher dimensions is the object of future work.

## AUTHOR CONTRIBUTIONS

J. S. Bryan IV wrote the manuscript and carried out all derivations, coding, and inference. S. Pressé conceived the project and oversaw all aspects of the project.

## ACKNOWLEDGMENTS

We would like to thank the Schuler lab at the Univ. of Zürich for providing us with previously published data. Steve Pressé acknowledges support from the NIH (grant No. R01GM134426 and R01GM130745) and NSF (Award no. 1719537).

## SUPPLEMENTARY MATERIAL

An online supplement to this article can be found by visiting BJ Online at http://www.biophysj.org.

## SUPPLEMENTARY INFORMATION

### S0.5 Derivation of the likelihood

Here we derive the likelihood distribution for a our photon measurements given a particle position. One important consequence of our Ito approximation, equation 5, is that our likelihood for single photon data will be equivalent to our likelihood for binned data. That is to say that neither photon arrival times nor ordering of photon colors within a time window provide any additional information about the particle position. We will start this section by deriving the likelihood for single photon measurements. After showing that the single photon measurements contain no additional information compared to binned photon measurements, we then derive the likelihood for binned photons (equations 6 and 7) that we use throughout this work.

We first write out the probability of collecting *J* photons with photon arrival times, ***T***, and photon colors, ***ϕ***, within a time window,

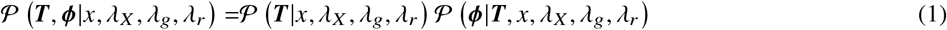

where, for simplicity, in our derivation we ignore artifacts such as crosstalk and detector efficiency. The time between photon arrivals will be exponentially distributed according to the excitation rate, *λ*_*X*_, and the background rates, *λ*_*g*_ and *λ*_*r*_. The probability of the photon arrival times, ***T*** is the probability of the *J* inter-photon times multiplied by the probability of no photon following the *J*th photon

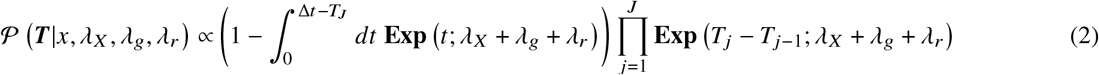

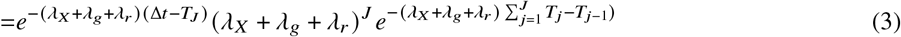

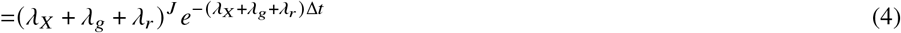

where in our derivation we used *T*_0_ = 0. The probability over the photon colors is the product of the probabilities over each individual photon given by the rates and the FRET efficiency

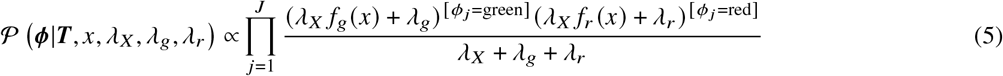

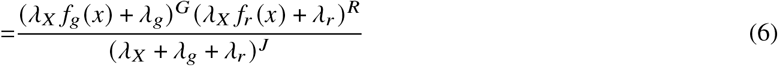

where *f*_*g*_ (*x*) = 1 − FRET (*x*), *f*_*r*_ (*x*) = FRET (*x*), [*x* = *y*] is the Iverson bracket (which is equal to 1 if *x* = *y* and 0 otherwise), and *R* and *G* are the total number of observed red and green photons. Putting this all together yields a distribution which has no dependency on individual photon arrival times nor photon color order,

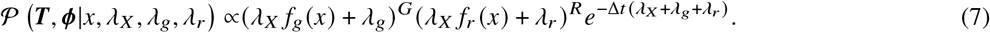

Since the likelihood depends neither on individual photon arrival times nor on photon color ordering, we lose no generality by rewriting our likelihood solely in terms of the number of measured photons within a time bin.

We now derive the likelihood for measuring *R* red photons and *G* green photons in a time window. The probability of collecting *G* green photons and *R* red photons in a time window is the probability of collecting *J* = *G* + *R* photons multiplied by the probability that *R* of the photons are red

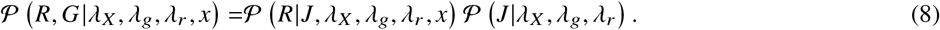

The probability of collecting *J* photons in a time window is Poisson distributed according to the rates,

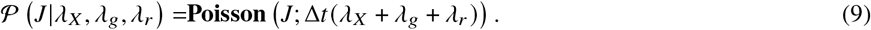

The probability that *R* photons are red is a binomial distribution with weight given by the relative rates of red and green photons

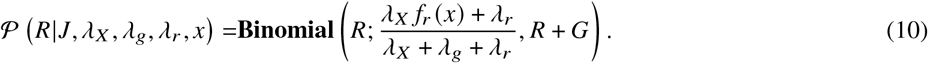

All together this yields

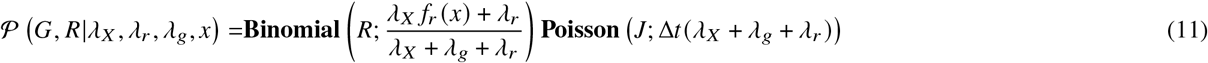

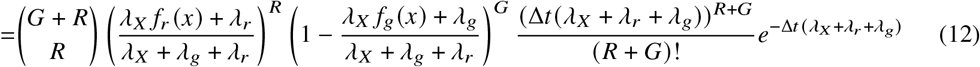

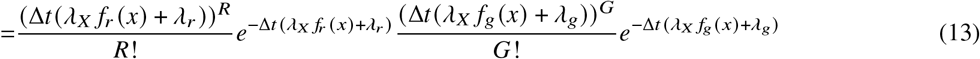

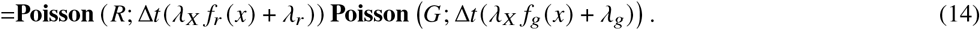

This is the likelihood (equations 6 and 7) that we use throughout this work.

### S0.6 Conditional probabilities

Here we derive the conditional probabilities used in the Gibbs sampling algorithm of section 0.3.

#### S0.6.1 Positions

The distribution for the positions is the product of the likelihood (equations 6 and 7), the dynamics model (equation 5), and the prior on the initial position (equation 16)

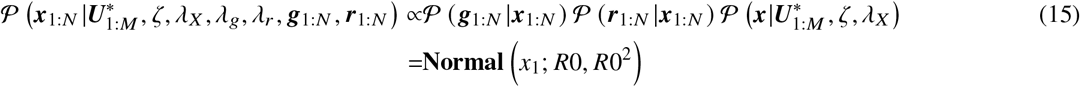

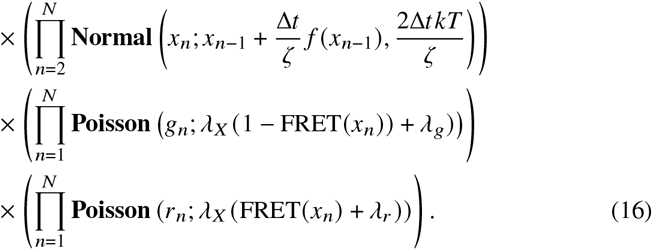

To sample from this distribution we may sample each *x*_*n*_ individually using a Metropolis Hastings (22) step. Separating equation 16 into conditional distributions on each position yields three equations: a conditional posterior on *x*_1_:

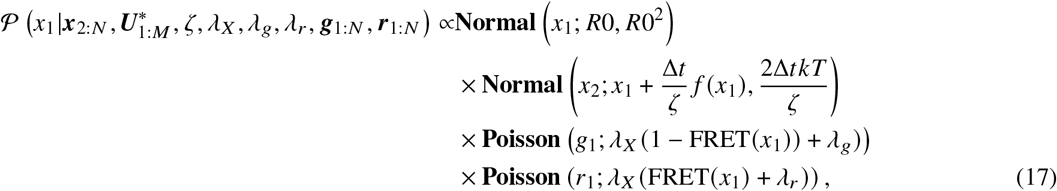

an equation for each *x*_*n*_ from time levels 2 to *N* − 1:

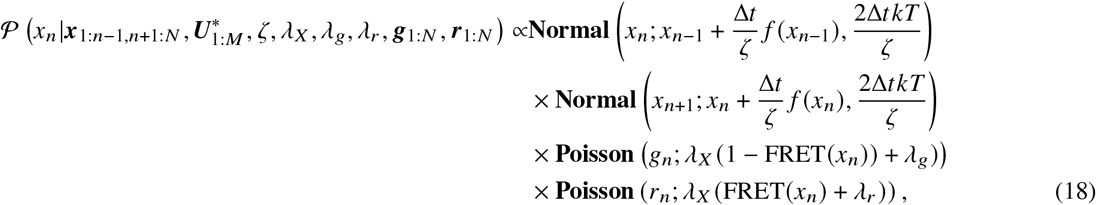

and an equation for the last position, *x*_*N*_,

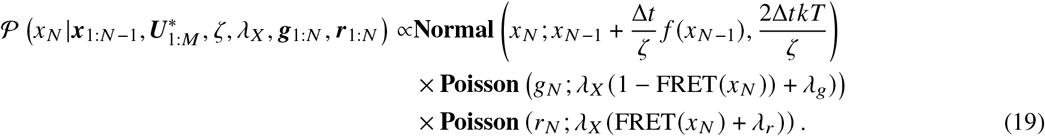

### S0.7 Potential

The conditional distribution for the potential is the product of the dynamics model (equation 5) and the prior on the potential (equation 17)

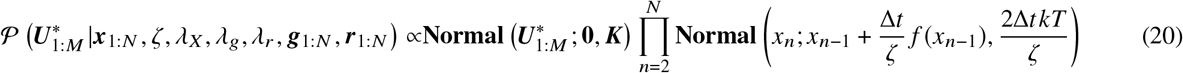

which can be simplified to (31, 32)

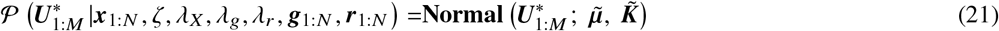

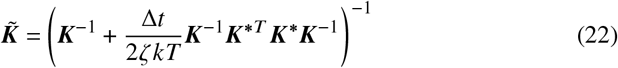

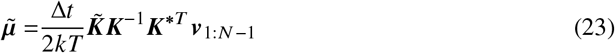

where ***K*** is the kernel matrix (covariance matrix) between all 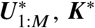 is the covariance between the potential at 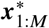 and the force at ***x***_1:*N*_ with elements 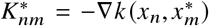, and _1:*N* −1_ are the velocities at each time level with elements ***v***_*n*_ = (*x*_*n* + 1_ − *x*_*n*_) /Δ*t*. As the final distribution for 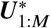 is Gaussian, we may directly sample 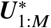 from the posterior without invoking Metropolis Hastings (31).

### S0.8 Photon rates

The conditional distribution on the excitation rate is the product of the likelihood (equations 7 and 6) and the prior on excitation rate (equation 12)

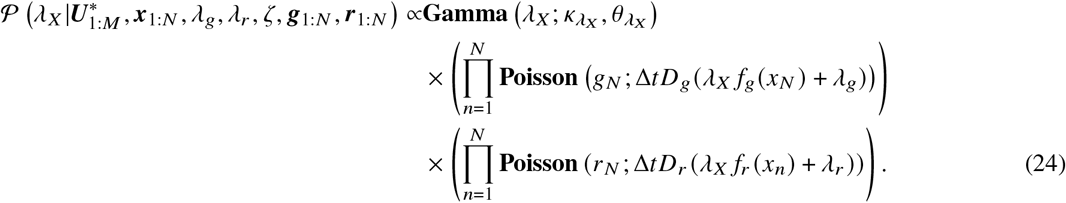

The distribution for the background rates is constructed in an identical manner except for the prior for which we replace the first term on the right hand side of 24 with equation 13 for *λ*_*r*_ and 14 for *λ*_*g*_. To sample from from either distribution we use Metropolis Hastings by proposing a sample at each iteration of the Gibbs sampler and accepting or rejecting based on the relative probabilities of the proposed variable and the old sample (22).

### S0.9 Friction coefficient

The conditional distribution over the friction coefficient is the product of the dynamics model (equation 5) and the prior on friction (equation 15)

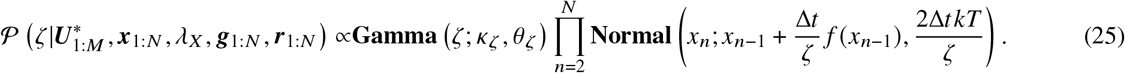

To sample from this distribution we use a Metropolis Hastings step by proposing a sample at each iteration of the Gibbs sampler and accepting or rejecting based on the relative probabilities of the proposed variable and the old sample (22).

### S0.10 Bayesian Hidden Markov model

We compare the energy landscape learned using our method to an energy landscape learned using a Bayesian Hidden Markov Model (HMM) (22, 47). Here we will briefly describe the structure of our HMM algorithm, then explain how we can use our Bayesian HMM analysis results to infer potential energy landscapes.

Briefly, HMMs work by assuming that the system under consideration has a discrete number of states, *k* = 1, 2, …, *K* governed by a transition matrix, ***q*** = [*q*_*i j*_]_*K*×*K*_. At each time level *n*, the system’s state, *s*_*n*_, is conditioned on the state of the system at the previous time level, *s*_*n*−1_, given the transition matrix, ***q***,

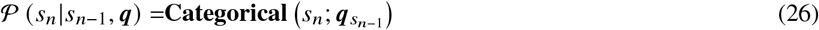

where 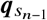 coincides with the row of ***q*** corresponding to *s*_*n*−1_. Put differently, the probability that *s*_*n*_ = *j* given that *s*_*n*−1_ = *i* is equal to *q*_*i j*_.

Each state, *k*, has its own pair distance, *r*_*k*_. At each time level, the measured number of photons is conditioned on the pair distance of the system’s state at that time level

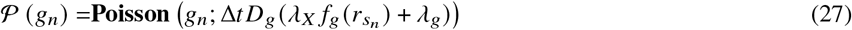

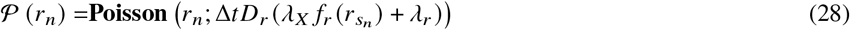

where *f*_*r*_(*x*) and *f*_*g*_(*x*) are the FRET rates, including crosstalk terms, defined by 9. Notice that this likelihood is equivalent to the SKIPPER-FRET likelihood, equations 6 and 7.

Working within the Bayesian paradigm, we place priors on all unknowns,

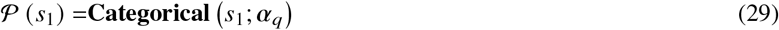

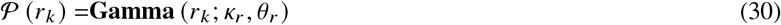

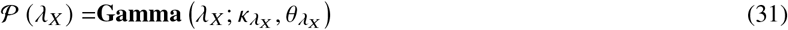

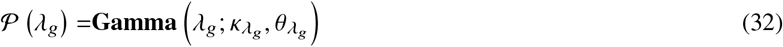

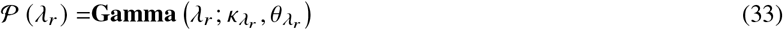

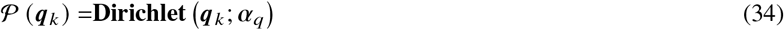

where **Dirichlet** (*α*_*q*_) is the Dirichlet distribution (22), conjugate to the Dirichlet dynamics model (equation 26) (22, 47). We choose our hyperparameters to be *α*_*q*_ = [1/ *K*, 1 /*K*, …, 1 /*K*], *κ*_*r*_ = 2, and *θ* _*r*_ = *R*_0_.

Equations (26) to (34) form a high dimensional posterior. We sample from our posterior using Gibbs sampling (22) and the forward filter-backward sampling algorithm (22). Once enough samples have been generated, we may choose to use the sample average to provide a point estimate for each variable.

In order to compare the HMM method to SKIPPER-FRET in the Results section, we used our HMM results to estimate the energy of each state. The energy of each state is calculated using the transition probability matrix, ***q***. We can find the energies from ***q*** by first calculating the equilibrium state probabilities, ***P***, defined as

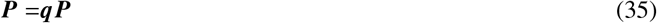

then equating ***P*** to the Boltzmann distribution

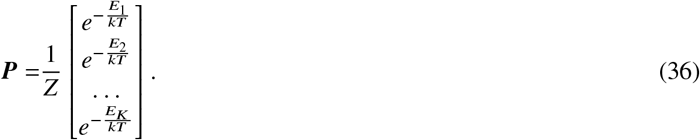

Together, equation 35 and 36 allow us to calculate the energy of each state in the HMM model.

#### S0.10.1 Barrier heights within HMM paradigm

Here we highlight how one would, if required, compute barrier heights within an HMM paradigm under two regimes: 1) features of the barrier are known; or 2) data are collected at different temperatures in addition to features of the barrier being known.

Here we focus on the first as it is of greater interest to experiments on biomolecules operating under one set of physiological temperatures.

**Figure S1:**
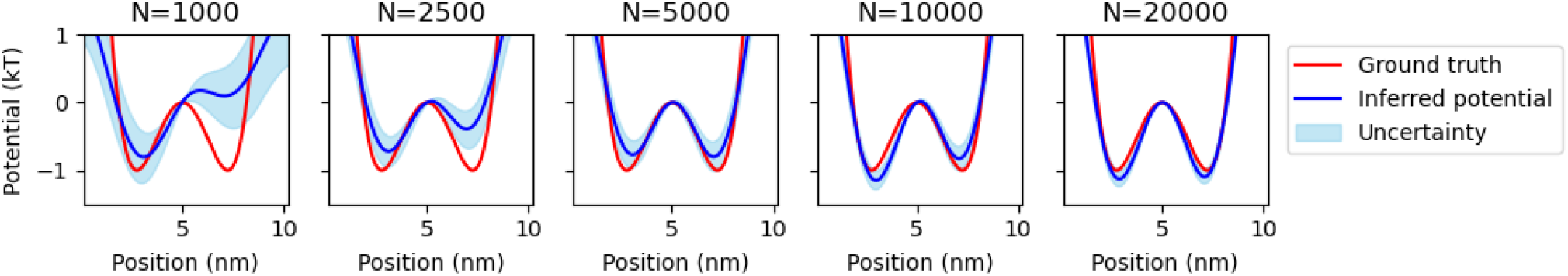
Robustness test with respect to number of data points. Each panel plots the potential inferred using SKIPPER-FRET (blue) against the ground truth potential energy landscape (red) for a given number of measurements, *N*, listed at the top.

In order to demonstrate that one could calculate barrier heights between states in the HMM model, we would first need to assume that the transition probability matrix, ***q***, is the solution to a master equation for a rate matrix, ***λ***,

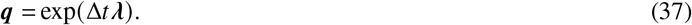

Solving for *λ* we get

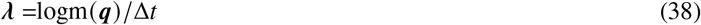

where logm is the matrix logarithm. Assuming that the wells representing each state can be approximated as harmonic oscillators, we can relate ***λ*** to barrier heights using Kramer’s rate equation (2, 20, 28)

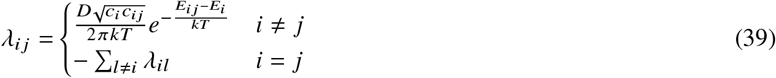

where *c*_*i*_ is the curvature of the well defining state *i, c*_*i j*_ is the curvature of the barrier between states *i* and *j, E*_*i j*_ is the energy of the barrier between states *i* and *j*, and *D* is a diffusion parameter dictating the rate of transitions in the absence of a barrier. Solving for the barrier heights we get

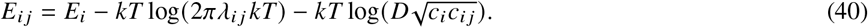

We note that using equation (40), we can only learn the energy of the barrier, *E*_*i j*_, if we know *D, c*_*i*_, and *c*_*i j*_. However, *D, c*_*i*_, and *c*_*i j*_ are internal parameters of the system which are not otherwise easy to deduce (28). In practice, bounds for barrier height are obtained by using additional approximations and an order of magnitude guess for unknown quantities (2, 28).

Thus we see that the inability to infer accurate potential energy barriers from a single data set without knowledge of hidden internal parameters is a major limitation of HMMs applied to smFRET data. By contrast, SKIPPER-FRET can learn barrier heights and friction coefficients from a single data set.

## S1 ROBUSTNESS WITH RESPECT TO NUMBER OF DATA

Here we test SKIPPER-FRET’s robustness with respect to the length of the data set. That is, we test how well the inferred potential energy landscape matches the ground truth given different number of time levels, *N*, available in the data. For our robustness test, we use the same simulated data as the first double well experiment (figures 3 and 4), but truncated at different values of *N*.

Figure S1 shows the results. As expected, when there are too few time levels for the pair distance to sample both wells, as is the case for *N* = 1000 and *N* = 2500 in figure S1, then SKIPPER-FRET cannot infer an accurate potential energy landscape due to missing data on the other well. When there is a sufficient number of time levels for the pair distance trajectory to sample both wells, as is the case for *N* ≥ 5000 in figure S1, then the form of the SKIPPER-FRET potential matches more closely with the ground truth potential energy landscape. Generally, figure S1 shows that the accuracy of SKIPPER-FRET increases with the number of time levels, and the uncertainty decreases with the number of data points. Note that the SKIPPER-FRET computation time increases linearly with the number of time levels.

Thus it is important to make sure that sufficient number of data are collected before applying SKIPPER-FRET. An ideal data set will have enough time for the pair distance to explore the whole space. For the purposes of this manuscript, we set *N* = 10000 for all data sets analyzed because this value gave us a great balance between accuracy and computation speed.

As it pertains to analysis of real experiments, of course, we can only ascertain the form of the potential for this regions ever visited.

## REFERENCES

1. Rob Phillips et al. Physical biology of the cell. GARLAND SCIENCE, 2012.

2. Benjamin Schuler, Everett A Lipman, and William A Eaton. “Probing the free-energy surface for protein folding with single-molecule fluorescence spectroscopy”. In: NATURE 419.6908 (2002), pp. 743–747.

3. Zhiqiang Yan and Jin Wang. “Funneled energy landscape unifies principles of protein binding and evolution”. In: PROCEEDINGS OF THE NATIONAL ACADEMY OF SCIENCES 117.44 (2020), pp. 27218–27223.

4. Corey Weistuch and Steve Pressé. “Spatiotemporal organization of catalysts driven by enhanced diffusion”. In: THE JOURNAL OF PHYSICAL CHEMISTRY B 122.21 (2017), pp. 5286–5290.

5. Dmitrii E Makarov. “Barrier Crossing Dynamics from Single-Molecule Measurements”. In: THE JOURNAL OF PHYSICAL CHEMISTRY B 125.10 (2021), pp. 2467–2476.

6. Pratyush Tiwary and Michele Parrinello. “From metadynamics to dynamics”. In: PHYSICAL REVIEW LETTERS 111.23 (2013), p. 230602.

7. Jiang Wang and Andrew L Ferguson. “Nonlinear reconstruction of single-molecule free-energy surfaces from univariate time series”. In: PHYSICAL REVIEW E 93.3 (2016), p. 032412.

8. Jin Wang and Gennady M Verkhivker. “Energy landscape theory, funnels, specificity, and optimal criterion of biomolecular binding”. In: PHYSICAL REVIEW LETTERS 90.18 (2003), p. 188101.

9. Xiakun Chu et al. “Quantifying the topography of the intrinsic energy landscape of flexible biomolecular recognition”. In: PROCEEDINGS OF THE NATIONAL ACADEMY OF SCIENCES 110.26 (2013), E2342–E2351.

10. Benjamin Schuler and William A Eaton. “Protein folding studied by single-molecule FRET”. In: CURRENT OPINION IN STRUCTURAL BIOLOGY 18.1 (2008), pp. 16–26.

11. Ken A Dill and Justin L MacCallum. “The protein-folding problem, 50 years on”. In: SCIENCE 338.6110 (2012), pp. 1042–1046.

12. Peter Hänggi, Peter Talkner, and Michal Borkovec. “Reaction-rate theory: fifty years after Kramers”. In: REVIEWS OF MODERN PHYSICS 62.2 (1990), p. 251.

13. Alexander M Berezhkovskii, Leonardo Dagdug, and Sergey M Bezrukov. “Mean direct-transit and looping times as functions of the potential shape”. In: THE JOURNAL OF PHYSICAL CHEMISTRY B 121.21 (2017), pp. 5455–5460.

14. Pavel F Bessarab, Valery M Uzdin, and Hannes Jónsson. “Potential energy surfaces and rates of spin transitions”. In: ZEITSCHRIFT FÜR PHYSIKALISCHE CHEMIE 227.11 (2013), pp. 1543–1557.

15. Hongyun Wang and George Oster. “Energy transduction in the F 1 motor of ATP synthase”. In: NATURE 396.6708 (1998), pp. 279–282.

16. Shoichi Toyabe, Hiroshi Ueno, and Eiro Muneyuki. “Recovery of state-specific potential of molecular motor from single-molecule trajectory”. In: EPL (EUROPHYSICS LETTERS) 97.4 (2012), p. 40004.

17. Robert M Clegg. “Fluorescence resonance energy transfer”. In: CURRENT OPINION IN BIOTECHNOLOGY 6.1 (1995), pp. 103–110.

18. Robert M Clegg. “The history of FRET”. In: REVIEWS IN FLUORESCENCE 2006. SPRINGER, 2006, pp. 1–45.

19. Eitan Lerner et al. “Toward dynamic structural biology: Two decades of single-molecule Förster resonance energy transfer”. In: SCIENCE 359.6373 (2018), eaan1133.

20. Stuart Lindsay. Introduction to nanoscience. OUP OXFORD, 2009.

21. Björn Hellenkamp et al. “Precision and accuracy of single-molecule FRET measurements—a multi-laboratory benchmark study”. In: NATURE METHODS 15.9 (2018), pp. 669–676.

22. Christopher M Bishop. Pattern recognition and machine learning. SPRINGER, 2006.

23. Steve Pressé, Julian Lee, and Ken A Dill. “Extracting conformational memory from single-molecule kinetic data”. In: THE JOURNAL OF PHYSICAL CHEMISTRY B 117.2 (2013), pp. 495–502.

24. Steve Pressé et al. “Single molecule conformational memory extraction: p5ab RNA hairpin”. In: THE JOURNAL OF PHYSICAL CHEMISTRY B 118.24 (2014), pp. 6597–6603.

25. Ioannis Sgouralis et al. “A bayesian nonparametric approach to single molecule forster resonance energy transfer”. In: THE JOURNAL OF PHYSICAL CHEMISTRY B 123.3 (2018), pp. 675–688.

26. Sean A McKinney, Chirlmin Joo, and Taekjip Ha. “Analysis of single-molecule FRET trajectories using hidden Markov modeling”. In: BIOPHYSICAL JOURNAL 91.5 (2006), pp. 1941–1951.

27. Frederick Reif. Fundamentals of statistical and thermal physics. WAVELAND PRESS, 2009.

28. Yongli Zhang, Junyi Jiao, and Aleksander A Rebane. “Hidden Markov modeling with detailed balance and its application to single protein folding”. In: BIOPHYSICAL JOURNAL 111.10 (2016), pp. 2110–2124.

29. Andrew Wilson and Hannes Nickisch. “Kernel interpolation for scalable structured Gaussian processes (KISS-GP)”. In: INTERNATIONAL CONFERENCE ON MACHINE LEARNING. PMLR. 2015, pp. 1775–1784.

30. Robert Zwanzig. Nonequilibrium statistical mechanics. OXFORD UNIVERSITY PRESS, 2001.

31. J Shepard Bryan IV et al. “Inferring potential landscapes from noisy trajectories of particles within an optical feedback trap”. In: ISCIENCE 25.9 (2022), p. 104731.

32. J Shepard Bryan IV, Ioannis Sgouralis, and Steve Pressé. “Inferring effective forces for Langevin dynamics using Gaussian processes”. In: THE JOURNAL OF CHEMICAL PHYSICS 152.12 (2020), p. 124106.

33. Christopher K Williams and Carl Edward Rasmussen. Gaussian processes for machine learning. Vol. 2. MIT PRESS CAMBRIDGE, MA, 2006.

34. Franziska Zosel et al. “Depletion interactions modulate the binding between disordered proteins in crowded environments”. In: PROCEEDINGS OF THE NATIONAL ACADEMY OF SCIENCES 117.24 (2020), pp. 13480–13489.

35. Flurin Sturzenegger et al. “Transition path times of coupled folding and binding reveal the formation of an encounter complex”. In: NATURE COMMUNICATIONS 9.1 (2018), pp. 1–11.

36. Franziska Zosel et al. “A proline switch explains kinetic heterogeneity in a coupled folding and binding reaction”. In: NATURE COMMUNICATIONS 9.1 (2018), pp. 1–10.

37. Ayush Saurabh et al. “Single photon smFRET. I. theory and conceptual basis”. In: BIORXIV (2022).

38. Ayush Saurabh et al. “Single photon smFRET. II. application to continuous illumination”. In: BIORXIV (2022).

39. Laura Baltierra Jasso, Michael Morten, and Steven William Magennis. “Sub-ensemble monitoring of DNA strand displacement using multiparameter single-molecule FRET”. In: CHEMPHYSCHEM 19.5 (2018), pp. 551–555.

40. Nolan Harris et al. “Is End-to-End Distance a Good Reaction Coordinate?” In: BIOPHYSICAL JOURNAL 96.3 (2009), 290a.

41. Narendar Kolimi et al. “Out-of-Equilibrium Biophysical Chemistry: The Case for Multidimensional, Integrated Single-Molecule Approaches”. In: THE JOURNAL OF PHYSICAL CHEMISTRY B 125.37 (2021), pp. 10404–10418.

42. Xinyu A Feng, Matthew F Poyton, and Taekjip Ha. “Multicolor single-molecule FRET for DNA and RNA processes”. In: CURRENT OPINION IN STRUCTURAL BIOLOGY 70 (2021), pp. 26–33.

43. Lin Wang and Weihong Tan. “Multicolor FRET silica nanoparticles by single wavelength excitation”. In: NANO LETTERS 6.1 (2006), pp. 84–88.

44. Janghyun Yoo et al. “Fast three-color single-molecule FRET using statistical inference”. In: NATURE COMMUNICATIONS 11.1 (2020), pp. 1–14.

45. Joseph P Torella et al. “Identifying molecular dynamics in single-molecule FRET experiments with burst variance analysis”. In: BIOPHYSICAL JOURNAL 100.6 (2011), pp. 1568–1577.

46. Tianyu Hu, Michael J Morten, and Steven W Magennis. “Conformational and migrational dynamics of slipped-strand DNA three-way junctions containing trinucleotide repeats”. In: NATURE COMMUNICATIONS 12.1 (2021), pp. 1–8.

47. J Shepard Bryan IV, Ioannis Sgouralis, and Steve Pressé. “Diffraction-limited molecular cluster quantification with Bayesian nonparametrics”. In: NATURE COMPUTATIONAL SCIENCE 2.2 (2022), pp. 102–111.

